# Within-host dynamics explain patterns of antibiotic resistance in commensal bacteria

**DOI:** 10.1101/217232

**Authors:** Nicholas G. Davies, Stefan Flasche, Mark Jit, Katherine E. Atkins

**Affiliations:** Centre for Mathematical Modelling of Infectious Diseases, London School of Hygiene and Tropical Medicine, London WC1E 7HT, UK; Department for Infectious Disease Epidemiology, Faculty of Epidemiology and Population Health, London School of Hygiene and Tropical Medicine, London WC1E 7HT, UK; Modelling and Economics Unit, Public Health England, London SE1 8UG, UK

## Abstract

The spread of antibiotic resistance, a major threat to human health, is poorly understood. Empirically, resistant strains gradually increase in prevalence as antibiotic consumption increases, but current mathematical models predict a sharp transition between full sensitivity and full resistance. In other words, we do not understand what drives persistent coexistence between resistant and sensitive strains of disease-causing bacteria in host populations. Without knowing what drives patterns of resistance, we cannot accurately predict the impact of potential strategies for managing resistance. Here, we show that within-host dynamics—bacterial growth, strain competition, and host immune responses—promote frequency-dependent selection for resistant strains, explaining patterns of resistance at the population level. By capturing these processes in a parsimonious mathematical framework, we resolve a long-standing conflict between theory and observation. Our models capture widespread coexistence for multiple bacteria-drug combinations across 30 European countries and explain associations between carriage prevalence and resistance prevalence among bacterial subtypes. A mechanistic understanding of resistance evolution is needed to accurately forecast the impact and effectiveness of resistance-management strategies.

---

Despite the global public health threat of antibiotic resistance^1, 2^, we currently lack a mechanistic understanding of how resistance spreads in human populations^3^. This gap in our knowledge is reflected in the inability of mathematical models to explain why resistance prevalence gradually increases as we move from countries with low antibiotic consumption to those with high antibiotic consumption^3–8^ (Fig. 1A). Although the mechanism driving this pattern seems intuitively obvious—greater antibiotic consumption selects for more resistance—the simplest models of disease transmission (Fig. 1B) are unable to capture the gradual rise that is observed. Instead, these models predict competitive exclusion^9^—that is, that above a particular level of antibiotic consumption, resistant strains fully outcompete sensitive strains, while below this treatment-rate threshold, they are unable to emerge at all (Fig. 1C).

**Fig. 1.**
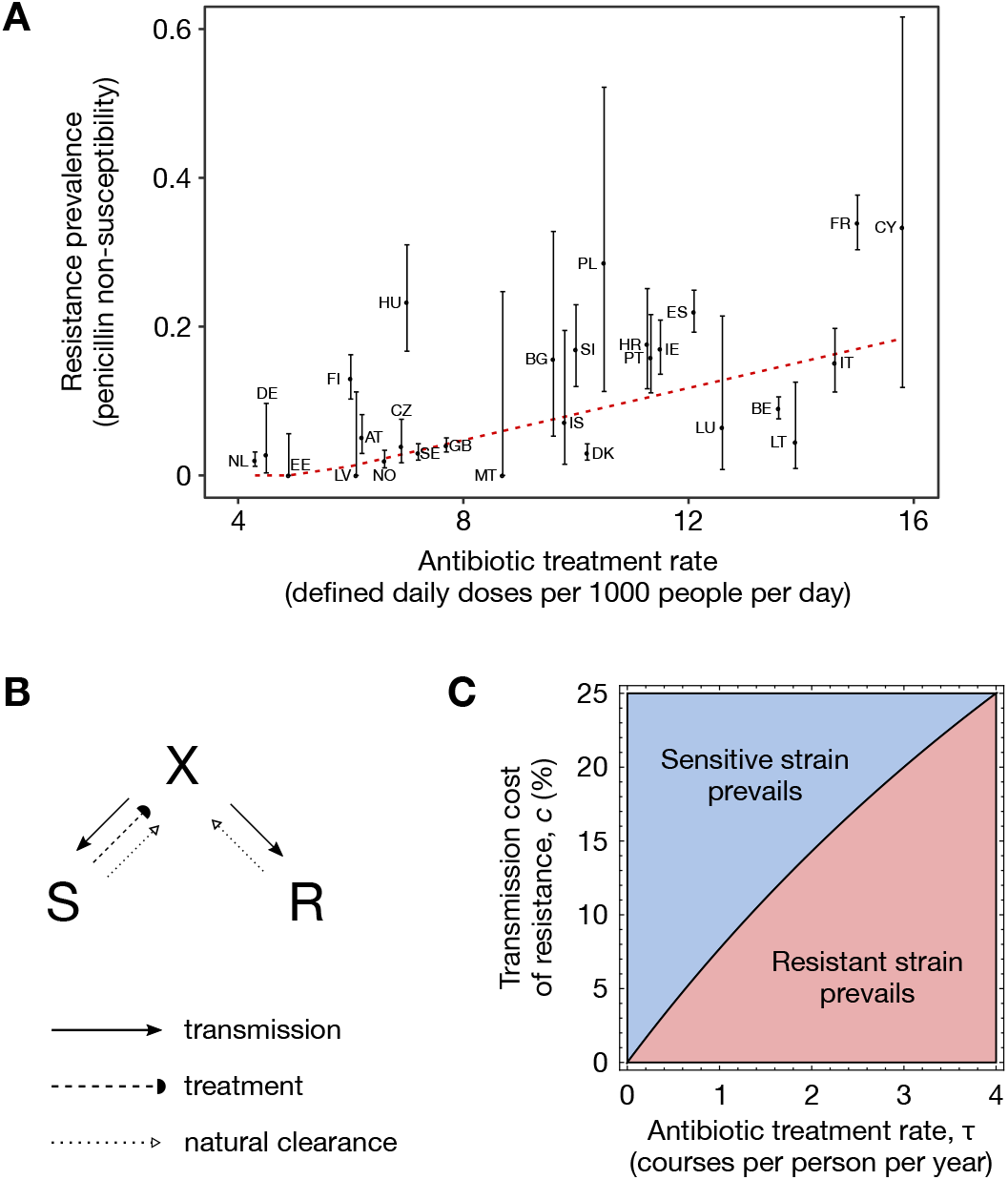
The problem of coexistence. **(A)** The prevalence of penicillin nonsusceptibility in invasive isolates of *S. pneumoniae* across 27 European countries illustrates widespread coexistence. This pattern of gradual increase of resistance with the antibiotic treatment rate holds across many pathogen-drug combinations^6–8^. **(B)** A simple two-strain single-carriage model defined by the system of ordinary differential equations 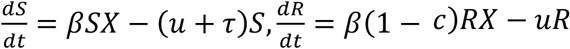, 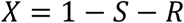, *X* = 1 – *S* – *R*, with *X* non-carriers, *S* sensitive-strain carriers, and *R* resistant-strain carriers. Here, *β* is the rate of transmission (solid arrows), *τ* is the rate of antibiotic treatment (dashed arrow), *u* is the rate of natural clearance (dotted arrows), and *c* is the transmission cost of resistance. **(C)** Contrary to observed patterns of coexistence, the single-carriage model predicts competitive exclusion, with the sensitive strain dominating the resistant strain if *τ* > *cu*/(1-*c*); here, *β* = 4 months^−1^, *u* = 1 months^−1^.

This theoretical prediction stands in stark contrast to empirical patterns of widespread coexistence between sensitive and resistant strains, a trend that holds across multiple bacteria-drug combinations^6–8^. While previous work has suggested that multiple strain carriage within individuals may promote limited amounts of coexistence^3, 10^, existing models cannot reproduce coexistence along the 4- to 20-fold range of treatment rates over which it is observed. Moreover, the reason why multiple carriage may promote limited coexistence is unclear. Without identifying an explicit, general mechanism for widespread coexistence, we will be unable to predict the likely impact of public health interventions for managing resistance.

Here, we show that within-host dynamics explain observed patterns of resistance in commensal bacteria. Specifically, we develop a suite of models that explicitly integrate within-host bacterial growth, strain competition, and bacterial subtype-specific host immune responses that, when calibrated to data across 30 European countries, provides a parsimonious and general explanation for empirical patterns of resistance in three commensal bacterial species, and also explains patterns of resistance among competing subtypes in the commensal bacterium *Streptococcus pneumoniae*.

## Results

### Existing models fail to capture widespread coexistence

We begin by analysing a standard model of resistant disease transmission developed by Lipsitch and colleagues^3, 10^. A central feature of this model is that hosts can become dual carriers—that is, carriers of both sensitive and resistant strains simultaneously—through sequential colonisation events (Fig. 2A). This model makes two key assumptions: (1) that dual carriage is balanced, with each strain carried in equal measure; and (2) that if a dual carrier is re-colonised, the incoming strain “knocks out” one of the two resident strains at random. Together, these assumptions preserve the crucial requirement of structural neutrality at the population level^10^, meaning that selection does not produce coexistence “for free” by artificially promoting transmission of rare strains. In this “within-host balancing” model, the amount of coexistence is governed by the parameter *k*, the efficiency of co-colonisation relative to single colonisation. While setting *k* = 0 eliminates dual carriage and recovers competitive exclusion, allowing dual carriage (0 < *k* ≤ 1) promotes limited coexistence (Fig. 2B). We calibrated this model to European data on penicillin use and penicillin nonsusceptibility among *S. pneumoniae (Methods*). Due to the limited range of coexistence predicted by the balancing model, we found that it cannot readily capture observed patterns of resistance^3, 5^ (Fig. 2C), even when co-colonisation is more efficient than single colonisation (k > 1; text S2).

**Fig. 2.**
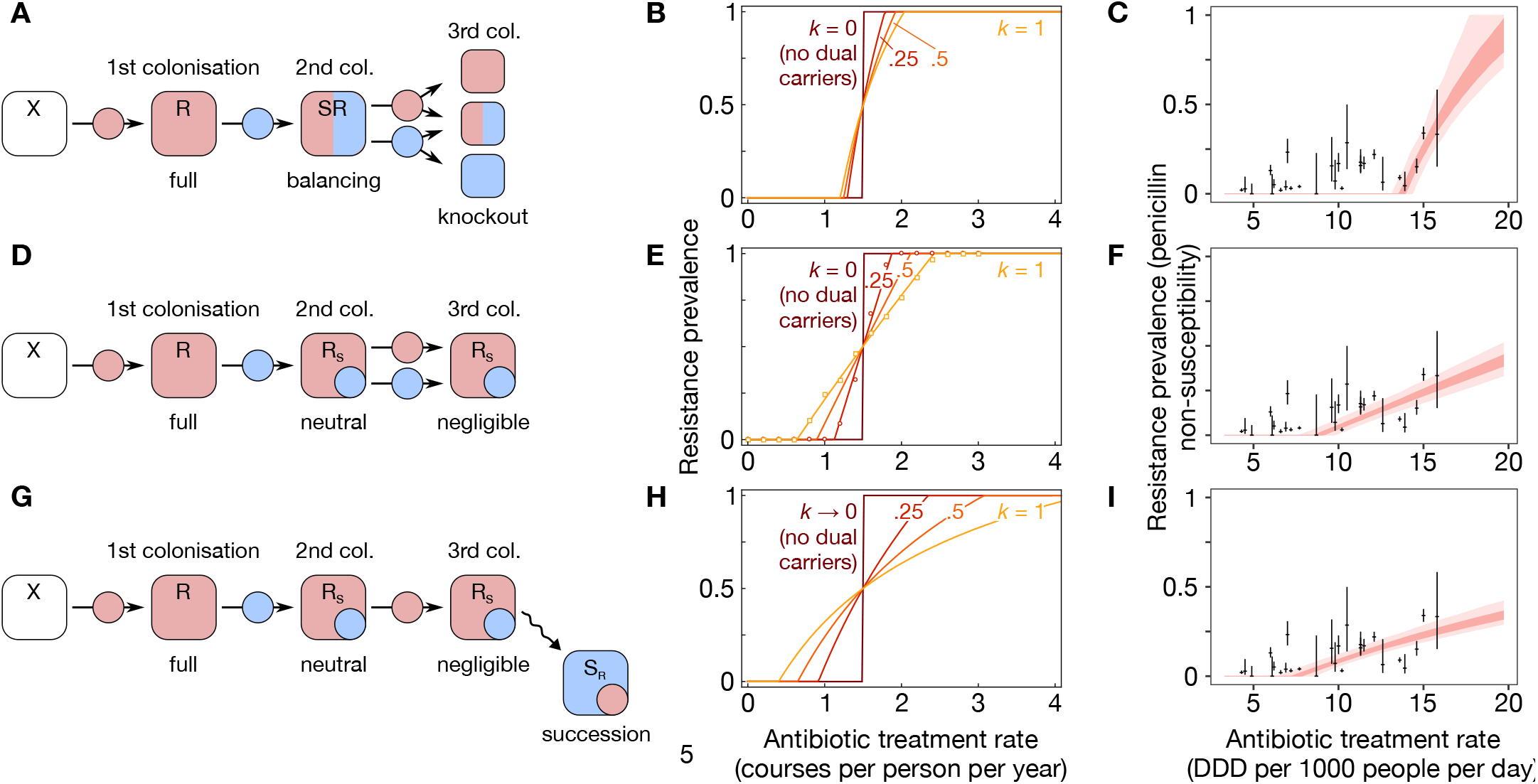
Three models of dual carriage. **(A)** The within-host balancing model^10^ extends the single-carriage model (Fig. 1B) by permitting simultaneous dual carriage. An example sequence of colonisation events illustrates equal co-colonisation and knockout. **(B)** Although coexistence increases with the efficiency of multiple carriage, *k*, the region of coexistence remains small for the within-host balancing model. **(C)** When calibrated to penicillin non-susceptibility in *S. pneumoniae* across European countries, the balancing model does not capture widespread coexistence. **(D)** The same example sequence of colonisation events as in panel **A** illustrates the absence of the balancing and knockout mechanisms in the within-host neutral model. **(E)** Integrating within-host neutrality expands the region of coexistence. Overlaid markers are results from our individual-based model with no limit to the number of co-colonisation states (see Methods). **(F)** Within-host neutrality yields a better fit to empirical data. **(G)** The same example sequence illustrates frequency-dependent selection acting on the sensitive strain when within-host competition is added to the within-host neutral model. **(H)** Within-host competition further increases coexistence and **(I)** the resulting model provides the best fit to the data (relative likelihood of 1.0 for the within-host competition model, 0.635 for the within-host neutral model, and 4.49×10^−5^ for the balancing model).

Although the balancing model preserves neutrality at the between-host level^10^, we argue that it violates the principle of structural neutrality at the within-host level. Specifically, a structurally-neutral model should not predict that balancing selection promotes stable coexistence between two strains if the strains are biologically identical^10^. The balancing model meets this requirement at the population level^3, 10^. However, at the within-host level, when an invading strain colonises a host already carrying a larger resident strain, the balancing model implicitly assumes that the invading strain multiplies within the host—at the expense of the resident strain—until both strains are present in equal frequency, even if the two strains are biologically identical. Moreover, if a dual carrier is recolonised, knockout immediately eliminates one of the host’s existing strains, an assumption which simplifies the mathematical model but which does not have a clear mechanistic explanation. In order to preserve neutral within-host dynamics, we developed a new model that relaxes these two non-neutral assumptions.

### Within-host neutrality captures widespread coexistence

In our novel “within-host neutral” model, which exhibits structural neutrality at all levels, hosts are assumed to have a fixed carrying capacity beyond which bacterial growth cannot be supported. If a host is already colonised, a new strain can invade, but rather than reaching the same frequency as the resident strain, we assume that the new strain does not increase in frequency after co-colonisation, since the host is already at carrying capacity (Fig. 2D). Strikingly, integrating within-host neutrality greatly promotes coexistence (Fig. 2E), allowing us to readily capture patterns of *S. pneumoniae* resistance across European countries (Fig. 2F). We originally developed this model as an individual-based simulation which allows us to track arbitrary within-host strain frequencies for each host, but the model can be approximated using a four-state system of differential equations, which simplifies analysis and model calibration (*Methods*).

It is generally agreed that antibiotic resistance is associated with a fitness cost^11, 12^, because in the absence of such a cost, resistant strains would fully outcompete sensitive strains. We have thus far parameterised this cost by assuming that resistant strains are associated with reduced transmission. Alternatively, it is possible to assume that resistant strains suffer reduced within-host growth^11, 12^, such that a resistant strain will, in the absence of antibiotic treatment, be outcompeted within the host by any sensitive strains (Fig. 2G). We captured this process of within-host competition using a five-state differential equation model based on our within-host neutral model (*Methods*). This addition further expands the parameter space over which coexistence is maintained (Fig. 2H), improving the model fit to patterns of *S. pneumoniae* penicillin non-susceptibility (Fig. 2I).

To verify the generality of our results, we re-parameterised the three models for two additional facultatively-pathogenic commensal bacteria, *Escherichia coli* and *Staphylococcus aureus* (*Methods*), and recalibrated the models to pan-European patterns of resistance against the most widely used antibiotics for all three species, which yielded a further four bacteria-drug combinations^6, 8^. Here, too, the empirical data are better captured by the within-host neutral models than by the balancing model (Fig. 3). Using the Akaike Information Criterion to select the most parsimonious model, we found that the within-host competition model has the most support across all bacteria-drug combinations (Figs. 2, 3).

**Fig. 3.**
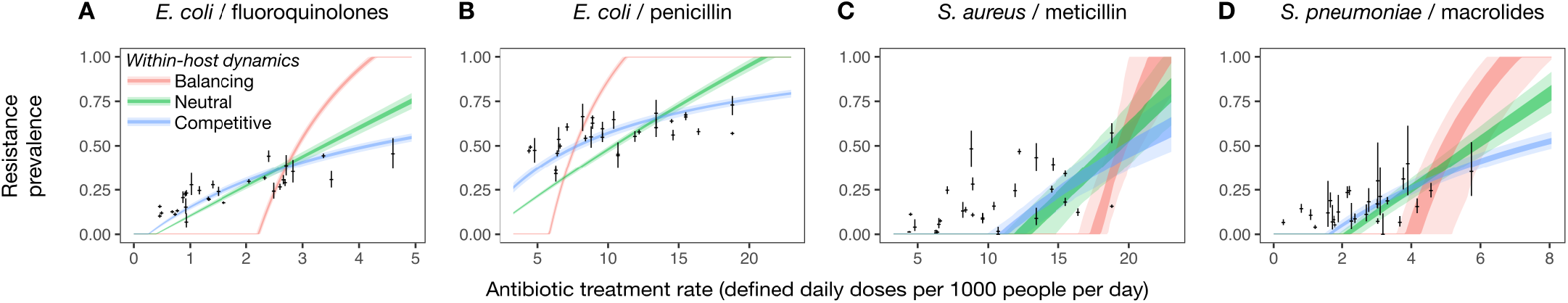
Within-host dynamics explain patterns of resistance across commensal bacteria. Within-host neutral models with (blue) or without (green) within-host competition capture persistent coexistence, while the balancing model (red) does not. Compared with the within-host neutral model with strain competition, the relative likelihood of the balancing and neutral models are, respectively, **(A)** 2.45×10^−127^, and 3.85×10^−15^; **(B)** 3.22×10^−163^ and 3.41×10^−28^; **(C)** 6.33×10^−4^ and 0.141; and **(D)** 6.57×10^−15^ and 2.28×10^−3^.

### Within-host dynamics promotes coexistence via frequency-dependent selection

Analysis of our model reveals that frequency-dependent selection^13^ is the mechanism through which within-host dynamics promote coexistence. That is, the within-host processes we have identified result in fitness advantages for either resistant or sensitive strains when those strains are rare. If resistant cells colonise a host already carrying a sensitive strain, and that host subsequently takes antibiotics, the sensitive strain is cleared and the resistant cells can grow to occupy the host’s now-vacated niche; by contrast, if resistant cells colonise a host already carrying a resistant strain, antibiotic treatment has no such effect. Hence, resistant cells benefit from a fitness advantage when sensitive-strain carriers are common. Within-host competition has a complementary effect, providing sensitive cells with a fitness advantage when resistant-strain carriers are common, because sensitive cells colonising a resistant-strain carrier can grow over time to become the dominant strain in the host. Either or both of these mechanisms can promote stable coexistence by equalising the fitness of resistant and sensitive strains at intermediate carriage frequencies.

Identifying the mechanism of frequency-dependent selection helps to explain why previous model predictions for resistance prevalence are sensitive to a narrow range of antibiotic treatment rates^3, 5, 10^, and hence have been unable to support coexistence over the wide range of treatment rates measured empirically. Specifically, the “balancing” and “knockout” assumptions of the within-host balancing model create a bias against coexistence in two ways. First, balanced carriage of co-colonising strains decreases rare strains’ frequency-dependent fitness benefit because it reduces the scope for rare strains to grow within dual carriers. Second, knockout reduces the prevalence of dual carriage by depleting strain diversity among hosts. Both effects reduce the strength of frequency-dependent selection for resistance, biasing models against coexistence.

### Patterns of coexistence among bacterial serotypes

Many bacterial species exhibit extensive diversity in the expression of capsular proteins exposed to host immune systems, which subdivides species into several distinct “serotypes” that, like resistant versus sensitive strains, are known to stably coexist in host populations^14, 15^. Host immune memory can promote coexistence between serotypes, because more common serotypes are more likely to provoke a host immune response owing to previous exposure, creating frequency-dependent selection for serotype diversity^14, 15^. However, similar considerations to those detailed above reveal that serotype diversity can also be promoted through a related mechanism that does not require host immune memory. In our within-host neutral model, coexistence is promoted by resistant strains multiplying to take the place of co-colonising sensitive strains that are cleared by antibiotic treatment. A similar mechanism might promote stable coexistence between commensal bacteria serotypes, so long as—when co-colonising a host— serotypes can be independently cleared by a host immune response, with any strains of the same serotype being cleared simultaneously. This would result in a fitness advantage for a strain co-colonising a host carrying a different serotype, promoting serotype diversity even in the absence of immune memory.

To test this hypothesis, we extended our individual-based model to track coexistence between five serotypes differing in their clearance rate, their transmission rate or their within-host growth rate. We found that independent clearance of serotypes from co-colonised hosts can promote stable coexistence between serotypes, even when serotypes exhibit intrinsic fitness differences (Fig. 4A). Unlike in previous models which have not explicitly tracked within-host dynamics^14, 15^, we find that acquired immunity is not needed to maintain limited amounts of serotype diversity. Moreover, we found that resistance prevalence varied across coexisting serotypes, with the direction of the trend in resistance prevalence among serotypes depending on whether the cost of resistance was parameterised as a transmission-rate cost or a growth-rate cost. When resistance is associated with a transmission-rate cost, resistance is more strongly selected in fitter serotypes, whether serotypes differ in fitness due to differences in duration of carriage^4^, transmissibility, or within-host growth (Fig. 4B. This reflects an empirical association between serotype fitness and resistance prevalence observed in *S. pneumoniae*^4^. By contrast, when resistance is associated with a within-host growth-rate cost (Fig. 4C), resistance is more strongly selected in less-fit serotypes (Fig. 4D). Thus, our model would predict that the association between resistance prevalence and serotype fitness may depend on the balance between these two putative costs of resistance.

**Fig. 4.**
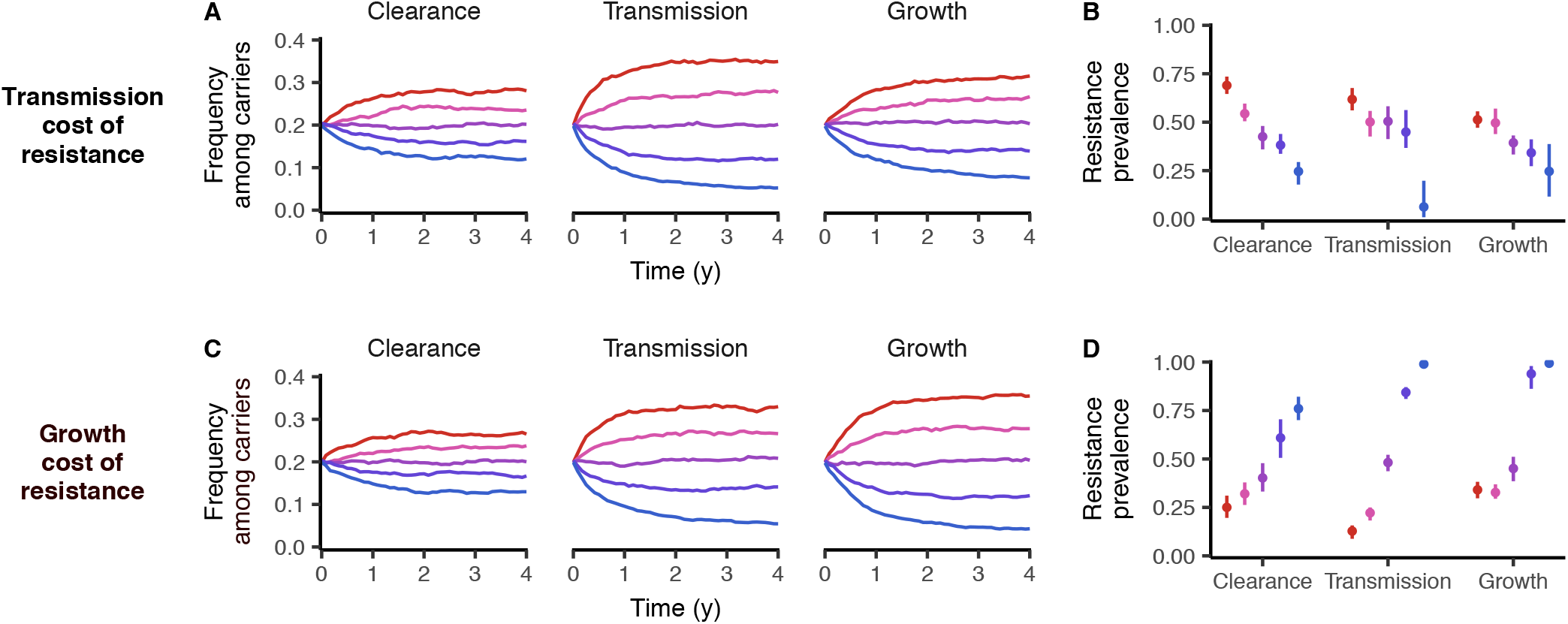
Within-host dynamics promote coexistence between serotypes, and intrinsic fitness differences between serotypes are correlated with resistance prevalence within serotypes. **(A)** In a model with five serotypes differing in various measures of intrinsic fitness, independent clearance of serotypes from hosts supports limited coexistence between serotypes even in the absence of immune memory. **(B)** When resistance carries a transmission-rate cost, fitter serotypes are more strongly selected for resistance. **(C, D)** When resistance carries a growth-rate cost, fitter serotypes are less strongly selected for resistance. Serotypes are assumed to differ in clearance rate (*u* = 0.96, 0.98, 1.0, 1.02, 1.04), transmission rate (β = 2.2, 2.1, 2.0, 1.9, 1.9), or within-host growth rates (growth rate penalties α = 0, 0.015, 0.03, 0.045, 0.06; see Methods). In each plot, the fittest serotype is shown in red. The mean and 95% interquantile range for the last 50 years of the 100-year simulation is shown. Resistance is assumed to be associated with either a 10% transmission-rate cost (A, B) or a growth-rate penalty of 0.025 (C, D).

### Resistance prevalence among pneumococcal serotypes

Finally, we used this proof of principle to evaluate carriage distribution and resistance prevalence in a model with 30 co-circulating *S. pneumoniae* serotypes parameterised with observed differences in duration of carriage. We found that independent clearance of serotypes alone was insufficient to support the high diversity of pneumococcal serotype carriage observed in human populations, with only five serotypes maintained (text S4). However, introducing serotype-specific immunity^14^ to our within-host neutral framework was sufficient to capture much of the observed pneumococcal diversity and patterns of resistance among pneumococcal serotypes (Fig. 5).

**Fig. 5.**
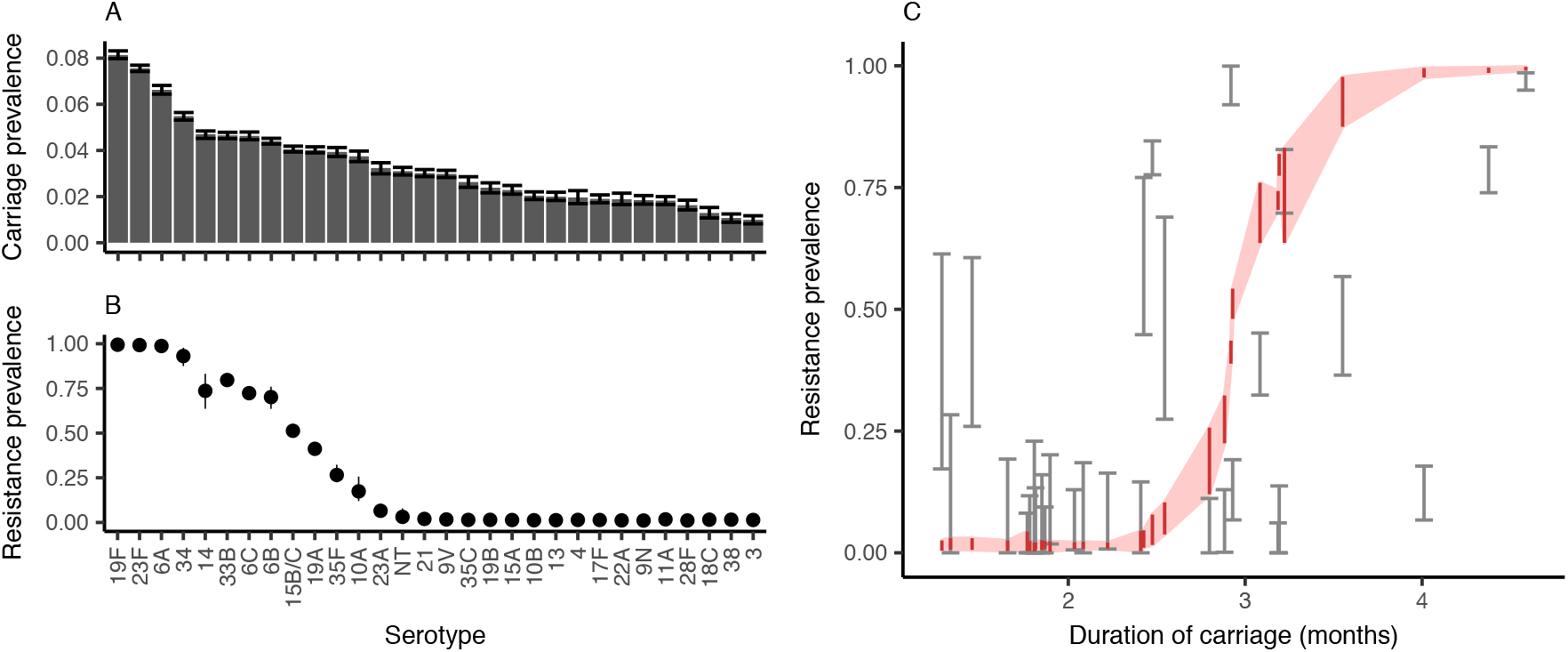
Resistance prevalence among pneumococcal serotypes. (A We parameterised our individual-based within-host neutral model to simulate a population in which 30 common pneumococcal serotypes circulate, using a previously-published data set^4, 31^ to assign measured durations of carriage to each serotype. Serotypes are otherwise undifferentiated and we assume no within-host competition. Incorporating a simple model of host immune memory (see *Methods*) recovers extensive diversity in **(A)** pneumococcal carriage and **(B)** resistance prevalence. **(C)** The positive relationship between duration of carriage and resistance prevalence observed empirically (black error bars, showing 95% confidence intervals for duration of carriage) is recovered in the model (red ribbon, showing 95% interquantile range for each serotype in the final 50 years of the 100-year simulation).

## Discussion

We have proposed a novel, parsimonious and pathogen-independent mathematical framework that, unlike previous models, captures empirical patterns of antibiotic resistance across countries differing in treatment rates and among serotypes. We argue that frequency-dependent selection drives these patterns of resistance and that coexistence of sensitive and resistant strains is promoted by integrating structural neutrality at the within-host level. Frequency-dependent selection has long been known to promote stable coexistence among animal, plant, and bacterial competitors^16–19^, and we show here that it could also have a pivotal role in maintaining coexistence between sensitive and resistant strains of commensal bacteria.

Although a number of alternative mechanisms that could explain coexistence between drug-sensitive and resistant pathogens have been proposed^3–5, 20–23^, some support only modest amounts of coexistence^3, 5^, while other proposed mechanisms may be difficult to generalize empirically, such as strongly age-assortative mixing^4^, independent mappings of balancing selection^4^, or specific immune responses to resistance-associated phenotypes^3, 21–23^. Our framework of within-host neutrality provides two advances on previous work: it harmonises pathogen dynamics occurring at the between-host and within-host levels, and it allows us to better and more parsimoniously capture observed patterns of resistance prevalence across a range of important bacteria-drug combinations.

Dual carriage of resistant and sensitive strains, a crucial factor for coexistence, is generated in our models via sequential colonisation. The empirical prevalence of dual carriage is not well known, but a study of *S. aureus* carriage in children found 21% of hosts carried both resistant and sensitive strains^24^ and—although resistance phenotypes were not measured—other studies have found up to 48% multiple carriage of genetically-distinct *S. pneumoniae* strains^25, 26^ and 42% multiple carriage of virulent *E. coli* strains^27^. These studies find that carriage is typically dominated by one strain with other strains carried at low frequency, consistent with our model’s carrying-capacity assumptions. Dual carriage could also occur through *de novo* mutation, which is likely to be especially important for long-lived chronic infections such as colonisations of the cystic fibrosis lung by *Pseudomonas aeruginosa*^28, 29^. While it is possible that coexistence is maintained by forces additional to dual carriage, we suggest that any model incorporating dual carriage should observe within-host neutrality to avoid biasing against coexistence.

Antibiotic resistance is one of the foremost threats to human health, and combating this threat will require the global deployment of coordinated interventions^1, 2^. Mathematical models of disease transmission will play a crucial role in this endeavour, because they can explicitly integrate the mechanisms that drive resistance evolution in a population-level framework and allow us to quantify long-term trends as well as the likely impact of any large-scale interventions for reducing resistance^30^. Providing a framework in which to answer public health questions will require a balance between mathematical tractability and identifiability of the models on one side, and necessary complexity on the other; building on the simple models proposed here will help to establish that balance. If mathematical models are able to incorporate a truly mechanistic understanding of resistance, they will be better able to explain common patterns of resistance and to accurately predict the effect of interventions at a national and global level^30^. With growing calls for an integrated and multifaceted approach to the problem of antimicrobial resistance^1, 2^, a new generation of mechanistic mathematical models will be uniquely placed to support the evidence-based adoption of impactful and cost-effective strategies.

## Acknowledgements

We thank M. Davies for assistance. NGD, MJ and KEA were funded by the National Institute for Health Research Health Protection Research Unit (NIHR HPRU) in Immunisation at the London School of Hygiene and Tropical Medicine in partnership with Public Health England (PHE. The views expressed are those of the authors and not necessarily those of the NHS, the NIHR, the Department of Health or PHE. The authors declare no competing financial interests. NGD, SF, MJ and KEA conceived the study; NGD performed the analyses; NGD and KEA drafted the manuscript, which all authors revised.

## Supplementary Materials

### Materials and Methods

#### Systems of ordinary differential equations

In the main text, we discuss four epidemiological models of antibiotic resistance: a single-carriage model, and three multiple-carriage models (the within-host balancing model, the within-host neutral model, and the within-host competition model). These models can be analysed using the systems of differential equations presented in this section, or using an alternative stochastic individual-based model formulation which is presented below (see *Individual-based mode*!). The ordinary differential equation models can simulate two strains at a time (sensitive and resistant), and are used for exposition of the key principles and for model calibration (Figs. 1–3). The individual-based models can simulate an arbitrary number of strains, and are used for validation of the ordinary differential equation models (Fig. 2) and for analysing serotype dynamics (Figs. 4 and 5).

*Single-carriage model* — The single-carriage model is given by

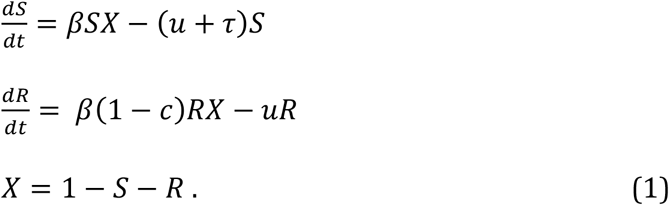

Here, *S* = *S*(*t*) is the fraction of sensitive strain carriers in the population; *R* = *R*(*t*) is the fraction of resistant strain carriers in the population; and *X* = *X*(*t*) is the fraction of non-carriers. The parameter β is the person-to-person transmission rate of the sensitive strain, *u* is the natural clearance rate, τ is the per capita rate of antibiotic treatment, and *c* is the transmission cost of resistance. Here, the resistance prevalence is ρ = *R*/(1–*X*).

*Within-host balancing model* — Following previous work^3, 10^, the balancing model is given by

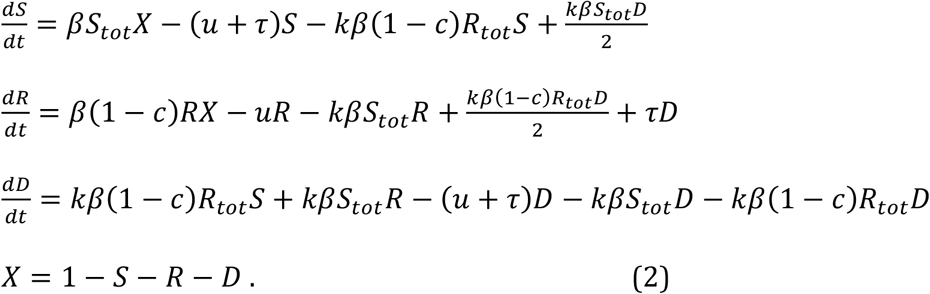

Here, the additional compartment *D* = *D*(*t*) is the fraction of dual carriers in the population (*SR* carriers in Fig. 2A of the main text), *k* is the efficiency of multiple colonisation relative to single colonisation, and *S*_tot_ = *S* + *D*/2 and *R*_tot_ = *R* + *D*/2 give the effective population burden of sensitive- and resistant-strain colonisation, respectively. The resistance prevalence is ρ = *R*_tot_/(1–*X*).

*Within-host neutrality model* — The novel neutral model is given by

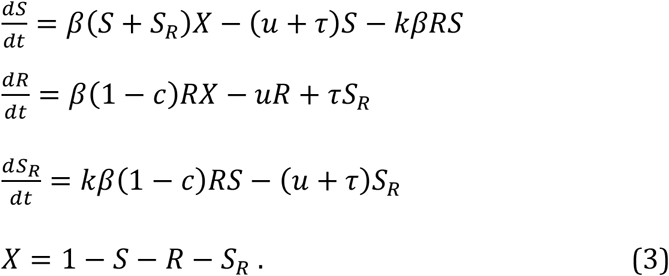

Here, the compartment *S_R_* = *S_R_*(*t*) captures the fraction of the population predominantly colonised with sensitive bacteria, but also carrying a small amount of resistant bacteria that are carried in insufficient quantity to transmit, and *S* + *S_R_* gives the effective population burden of sensitive-strain colonisation. The corresponding *R_S_* compartment is omitted as it does not impact upon the overall resistance prevalence in this model, ρ = *R*/(1–*X*); however, for tracking the overall prevalence of dual carriers (both *S_R_* and *R_S_*), the full neutral model can be recovered from equations 4 below by setting *b* = 0.

*Within-host competition model* — The neutral model with within-host competition is given by

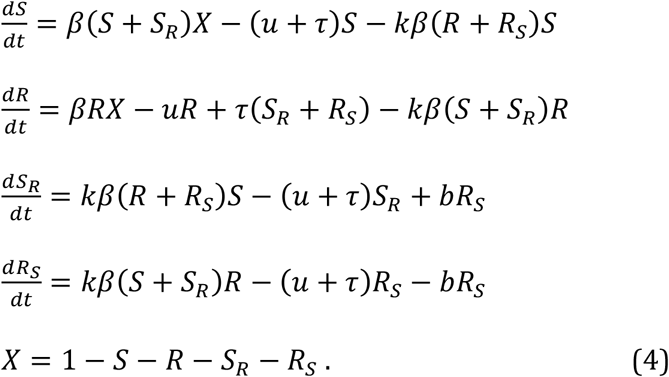

Here, the *R_S_* compartment captures dual carriers predominantly colonised with resistant bacteria plus a small amount of sensitive bacteria, and *b* captures the within-host benefit of sensitivity, *i.e*. the rate at which *R_S_* carriers become *S_R_* carriers. The resistance prevalence is ρ = (*R* + *R_S_*) /(1–*X*).

*Ordinary differential equation solutions* — All models are solved by setting singlecarriage compartments (S and R) equal to 0.001 and all dual-carriage compartments to 0, then integrating the systems of ordinary differential equations numerically in C++ using the Runge-Kutta Dorman-Prince method until they reach equilibrium.

#### Individual-based model

The individual-based model follows the same logic as the ordinary differential equation models, except it can simulate more than two strains simultaneously and within-host dynamics are simulated more explicitly.

In the individual-based model, hosts are indexed by *i* ∈ 1… *N*, where *N* is the population size, and there are *M* strains indexed by *j* ∈ 1… *M*. Each host is characterised by a vector (*f*_*i*,1_, *f*_*i*,2_,…, *f_i,M_*) giving the load of each strain colonising the host, where 0 ≤ *f_i,j_* ≤ 1 for all *i, j* and the host’s total bacterial load is *f_i_* = Σ_*j*_ *f_i,j_*. The force of infection of strain _*j*_ is *λ_j_* = (β/*N*) Σ_*i*_ *q_i_* (1–*c_j_*) *f_i,j_*, where β is the basic transmission rate, *c_j_* is the transmission cost associated with strain *j, q_i_* = 0 if *f_i_* = 0, and *q_i_* = 1/*f_i_* if *f_i_* > 0; that is, hosts with no bacterial load are not infectious, but all hosts with any nonzero bacterial load are assumed to be equally infectious, aside from differences in the transmission cost associated with each of their carried strains. Henceforth, we omit host subscripts *i* to focus on a single host.

The model proceeds in discrete time steps. Each time step, hosts may or may not experience a series of “events” (colonisation, treatment, or natural clearance) and then their strain carriage is “updated” according to a series of rules. For a time step Δ*t*, an event which occurs at rate *r* over the whole population is assumed to occur to each host with probability *r*Δ*t*. Colonisation events occur at rate *λ_j_* for each strain *j* if a host is uncolonised and rate *kλ_j_* if a host is already colonised. When a host is colonised by strain *j*, the host’s carriage of that strain is increased by the “germ size” *ι*, where *ι* ≪ 1, so we have 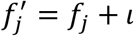, *i.e*. colonisation by strain *j* increases the host’s carriage of that strain by *ι* while leaving carriage of other strains unaffected. Additionally, hosts are colonised by importation of strains at a low rate ω = 1×10^−5^ to prevent stochastic elimination of strains at low carriage frequencies^14^. Treatment events occur at rate τ. When a host undergoes treatment, the host’s carriage of each strain becomes 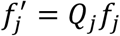 for all *j*, where *Q_j_* = 0 for sensitive strains and *Q_j_* = 1 for resistant strains, *i.e*. all sensitive strains are set to zero frequency. Clearance events occur at rate *u_s_* for each serotype s; each strain *j* is associated with a serotype *s_j_*, and there may be more than one strain with the same serotype. When a host undergoes natural clearance, the host’s carriage of each strain becomes 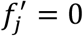 for all *j* for which *s_j_* = *s*, *i.e*. carriage of all strains of serotype *s* are set to zero while carriage of other strains is unaffected.

Once all “events” for a time step have been done, each host is “updated” according to the following rules. Under the within-host neutral model, total strain carriage for colonised hosts is always maintained at the carrying capacity of the host (i.e. *f* = Σ*f_j_* = 1). Specifically, a host’s bacterial load becomes 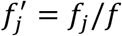 for all *j*. Under the within-host competition model, strain dynamics are modelled using an ordinary differential equation model of logistic growth and cell death; that is, we integrate from time *t* to time *t* + Δ*t* the differential equations defined by

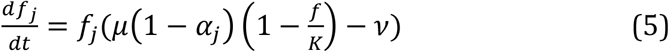

for all strains *j*, where μ is the basic growth rate, *α_j_* is the growth-rate penalty for strain j, *K* is the host density regulation constant, and ν is the cell death rate. For all simulations of the within-host competition model, we assume μ = 200, ν = 100, and *K* = 2, which maintains total strain carriage for each host at *f* = 1 when no strain has a growth-rate penalty. The model proceeds in discrete time steps of Δ*t* = 0.001 or Δ*t* = 0.005 months.

##### Host immune memory

When modelling the full range of pneumococcal serotypes (Fig. 5), we simulate host immune memory as follows. A host who has previously cleared a strain of serotype *s* (*i.e*. through natural clearance only, not treatment) cannot be reinfected by that serotype. However, hosts are randomly cleared of their immune memory at rate 1/*a*_max_, where *a*_max_ is the maximum host age simulated in the model, which approximates new hosts entering the population at birth at the same rate that hosts leave the population due to aging. We assume *a*_max_ = 60 months, reflecting the relative importance of pneumococcal transmission in hosts aged 5 years or under^3, 32^.

##### Model parameterisation and calibration

We calibrated the models using European Centre for Disease Prevention and Control data on penicillin consumption^6^ and the prevalence of penicillin non-susceptibility among *S. pneumoniae* invasive isolates^7^ from 2007 for 27 European countries. We focused on the year 2007 for this data set as criteria for penicillin non-susceptibility in *S. pneumoniae* were changed in some countries after this year, yielding inconsistencies in resistance data between countries^3, 33^. For calibration to all other bacteria-drug combinations, we used data for the same 27 countries collected in 2015, the most recent year for which data are available^6, 8^. Antibiotic consumption rates were given in defined daily doses (DDD) per thousand people; we converted these to overall treatment rates by assuming that 10 DDD comprise one treatment course for penicillin^4^ and fluoroquinolones, while 7 DDD comprise one treatment course for macrolides.

We used Markov chain Monte Carlo with differential evolution^34^ to calibrate the model output to empirical data. We assumed that the number of resistant isolates tested in a given country follows a binomial distribution with success probability drawn from a beta distribution, with the model-predicted resistance prevalence as the mode of the beta distribution. Setting the success probability as a random variable allowed us to account for between-country variation in resistance prevalence not captured by our dynamic model. The likelihood of the observed resistance prevalence for bacteria-antibiotic set *j* across all countries, ***d_j_***, given the model predicted resistance prevalence, *ρ_j_*, and prevalence of carriage, *Y_j_*, is

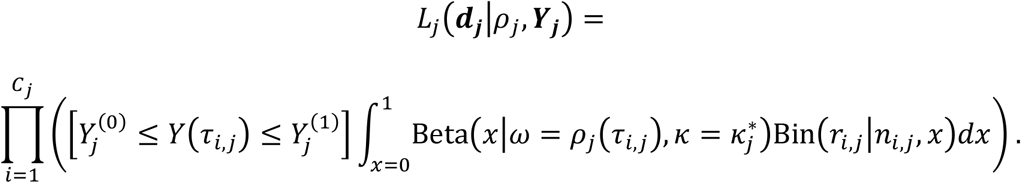

Here, 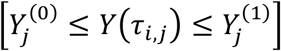 is 1 if *Y*(*τ_i,j_*) lies between 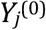 and 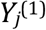, which are the lower and upper bounds of duration of carriage, respectively, and 0 otherwise; Beta(x|ω,κ) = x^α-1^(1-x)^β-1^Γ(α+β)/(Γ(α) Γ(β))|α = ω(κ-2) + 1, β = (1-ω) (κ-2)+1 is the beta distribution probability density with mode ω and concentration κ; and 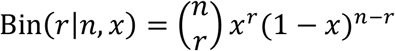 is the binomial distribution probability density. There are *C_j_* countries in the data set *j* labelled *i* = 1…*C_j_*; for a country *i* and bacteria-antibiotic set *j*, the measured antibiotic use is *τ_i,j_*, the number of samples taken of a particular bacteria is *n_i,j_*, and the number of those samples tested as non-susceptible to the particular antibiotic is *r_i,j_*. The model prediction for resistance prevalence given treatment rate τ is *ρ_j_*(τ), the model prediction for prevalence of carriage given treatment rate τ is *Y_j_*(τ), and the estimated concentration around the best-fit line, estimated from the data, is 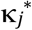.

To estimate the concentration 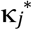 from the resistance prevalence data, we used a truncated line function *f*(*τ*) = max(0, *a* + *g*τ) as the modal antibiotic resistance prevalence and fixed 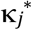 as the concentration. We calculated the maximum likelihood estimates of 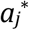, 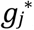, and 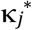 by using the Nelder-Mead algorithm^35^ to maximize the following likelihood function:

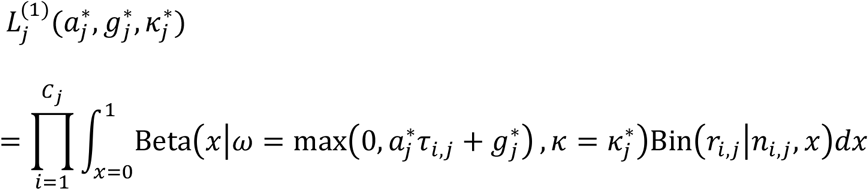

Priors for all model fitting were identified from the literature and were bacteria-specific (see below.

For illustrative purposes, the parameters *c* and *b* for Fig. 2B, E, H were chosen such that τ = 1.5 courses per person per year would lead to 50% resistance prevalence. This τ was chosen to roughly coincide with *S. pneumoniae* empirical data^6, 8^. Other parameters used were β = 4, *u* = 1 (ref. 3) and *k* as given in the figure.

###### Priors for duration of carriage

For *S. pneumoniae*, we assume an average duration of carriage of 1 month (ref. 3), giving a clearance rate *u* = 1 months^−1^. For *E. coli*, we assume that the average duration of carriage is 59 to 98 days^36, 37^, and accordingly set a uniform prior for *u* over the range 0.3–0.5 months^−1^. For *S. aureus*, we assume a duration of carriage of 3.3 months (estimated from ambulatory patients clearing MRSA^38^) to 5.9 months (ref. 39), and accordingly set a uniform prior for *u* over the range 0.17–0.3 months^−1^.

The transmission parameter, β, is constrained by the likelihood function (see below), so we set a uniform prior on β wide enough to overlap the full range of observed carriage prevalence, *Y*, given the range of clearance rates *u* and treatment rates *τ* (i.e. *Y* = 1 - (*u* + τ)/β for the sensitive strain alone, and *Y* = 1 - *u*/(β(1-*c*)) for the resistant strain alone). Specifically, for *S. pneumoniae*, we use the range 1–6; for *E. coli*, we use the range 1–10; and for *S. aureus*, we use the range 0.1–2.

###### Likelihood ranges for carriage prevalence

In the main text, we detail how the likelihood function used in model fitting constrains model output such that all countries must exhibit a prevalence of carriage *Y* such that *Y*^(0)^ ≤ *Y* ≤ *Y*^(1)^. For *S. pneumoniae*, we follow Colijn *et al*.^3^ in assuming 0.2 ≤ *Y* ≤ 0.8 in children. For *E. coli*, we restrict our attention to extraintestinal pathogenic *E. coli* (ExPEC), a subset of *E. coli* that is responsible for most invasive infections, and assume 0.499 ≤ *Y* ≤ 0.942, which is derived from 95% confidence intervals around the observed prevalence of carriage of ExPEC (9 of 12 healthy control subjects) from Martinez-Medina *et al*.^40^. Finally, for *S. aureus*, we adopt 0.2 ≤ *Y* ≤ 0.6 based on Bogaert *et al*.^32^, which is based on the observed range of *S. aureus* colonization in Dutch children, with a slightly wider range to capture more variation between European countries.

###### MCMC details

Following the differential evolution MCMC algorithm^34^, we run 10n chains, where *n* is the number of free parameters in the model, *i.e*. 30 for *S. pneumoniae* (for which carriage duration is fixed at 1 month) and 40 for *E. coli* and *S. aureus*, for which carriage duration is not fixed. The burn-in period lasts 1,000 iterations, after which 100,000 samples from the posterior are taken across all chains.

##### Sources for antibiotic consumption and resistance prevalence data

We used data from the European Centre for Disease Prevention and Control (ECDC and from the European Antimicrobial Resistance Surveillance Network (EARS-Net) to parameterize our models for the consumption of antibiotics and prevalence of antibiotic resistance in EU countries.

The ECDC provides information on antimicrobial consumption in defined daily doses (DDD) per 1000 inhabitants per day for participating countries, with antimicrobials broken down into categories as classified in the World Health Organisation’s (WHO Anatomical Therapeutic Chemical (ATC Classification System. Consumption is classified into primary care and hospital use, with the majority of use in primary care. We focus on primary care usage only, as hospital usage data is sparse and we are focusing on community-acquired bacterial carriage. The ECDC releases online publications summarizing consumption data (e.g. ref. 41) and maintains a queryable online database^6^.

EARS-Net, formerly EARSS, provides information on resistance levels as measured in invasive isolates of various bacterial pathogens isolated from blood and cerebrospinal fluid^7, 8^. We assume that invasive isolates are a sample of commensally-carried strains^4^. We summarize the data used and sources for the five data sets analysed (Table S1).

**Table S1.** Data sources for antibiotic consumption and antimicrobial resistance across five pathogen-drug combinations.

| Data set | Consumption:<br>*ATC code, sector, year, and source* | Resistance:<br>*Pathogen, resistance metric, year, and source* |
| --- | --- | --- |
| *Streptococcus pneumoniae* penicillin resistance, 2007 (Figs. 1–2, main text) | J01C (Beta-lactam antibacterials, penicillins), primary care, 2007 (ECDC 2017b, 2014<sup>†</sup>) | *S. pneumoniae*, percentage non-susceptible to penicillin, 2007 (EARSS 2007) |
| *Escherichia coli* fluoroquinolone resistance, 2015 (Fig. 3a, main text) | J01MA (Fluoroquinolones), primary care, 2015 (ECDC 2017b) | *E. coli*, percentage resistant to fluoroquinolones, 2015 (ECDC 2017a) |
| *E. coli* penicillin resistance, 2015 (Fig. 3b, main text) | J01C (Beta-lactam antibacterials, penicillins), primary care, 2015 (ECDC 2017b) | *E. coli*, percentage resistant to aminopenicillins, 2015 (ECDC 2017a) |
| *Staphylococcus aureus* meticillin resistance, 2015 (Fig. 3c, main text) | J01C (Beta-lactam antibacterials, penicillins), primary care, 2015 (ECDC 2017b) | *S. aureus*, percentage resistant to meticillin (MRSA), 2015 (ECDC 2017a) |
| *S. pneumoniae* macrolide resistance, 2015 (Fig. 3d, main text) | J01FA (Macrolides), primary care, 2015 (ECDC 2017b) | *S. pneumoniae*, percentage non-susceptible to macrolides, 2015 (ECDC 2017a) |

^†^Portugal recorded no penicillin consumption for 2007 in the online ECDC database^6^, but the 2011 ECDC report^41^ provides the corrected figure of 11.3 DDD per 1000 inhabitants per day for 2007.

### Supporting information

#### S1. Posterior distributions from model fitting

Here, we show the posterior distributions from model fitting from the main analysis (Fig. S1–S5).

**Figure S1.**
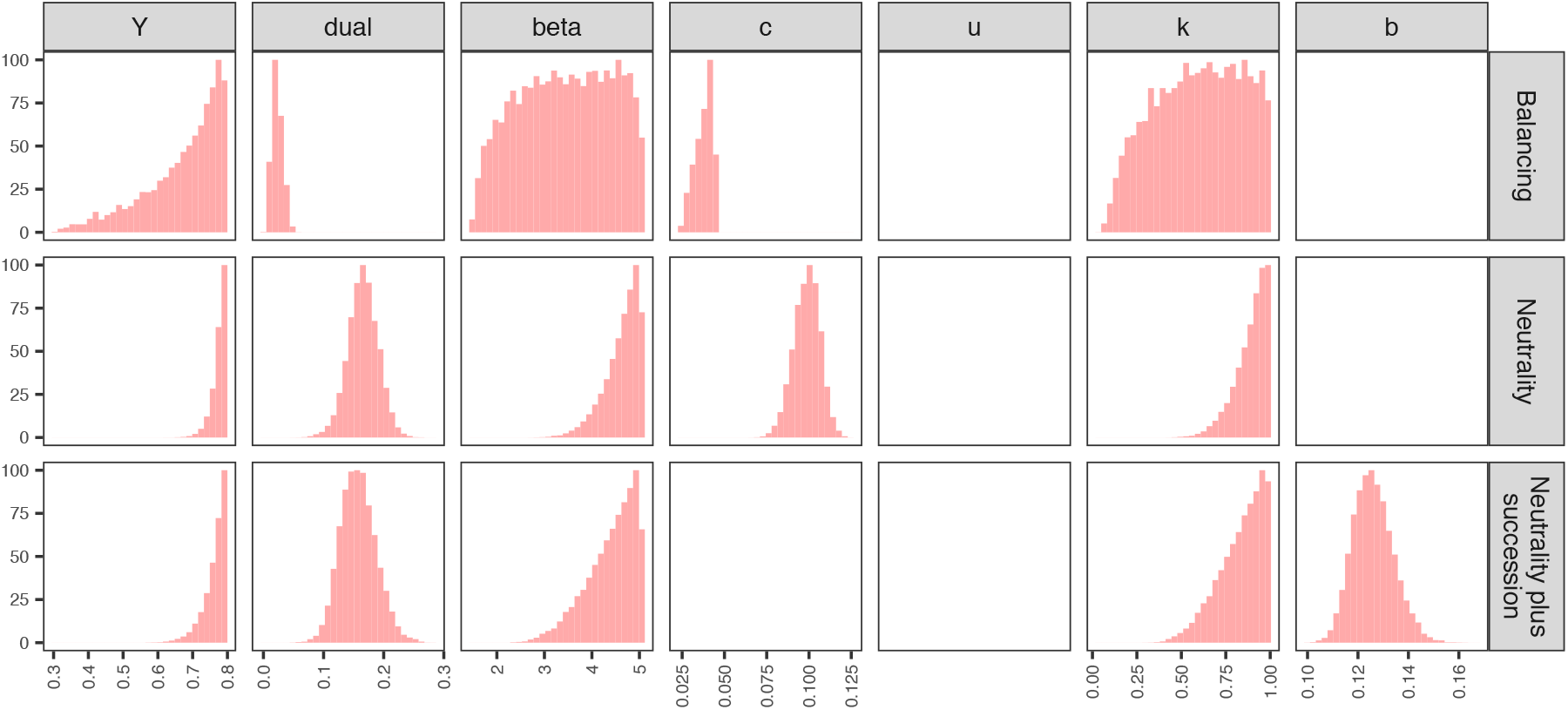
Posterior distribution for *S. pneumoniae* / penicillin, 2007. *Y* gives the overall prevalence of carriage (whether single or dual) in the population; *dual* gives the fraction of carriers who carry both sensitive and resistant strains; *beta* gives the transmission parameter; *c* gives the transmission cost of resistance; *u* gives the clearance rate (if the clearance rate is subject to fitting; for *S. pneumoniae, u* = 1); *k* gives the efficiency of recolonisation, and *b* gives the within-host rate of succession (see *Methods*, main text, for details).

**Figure S2.**
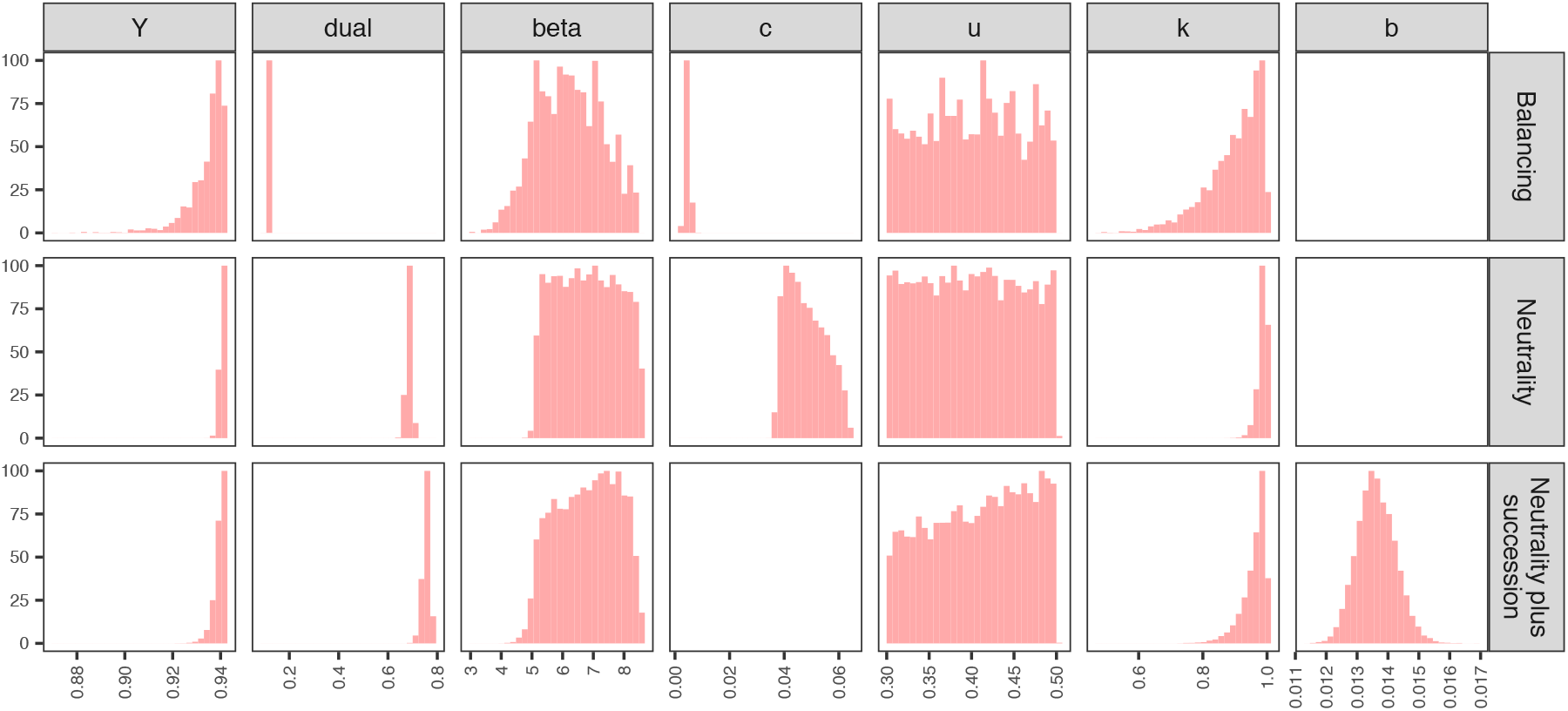
Posterior distribution for *E. coli* / fluoroquinolones, 2015. See Fig. S1 for details.

**Figure S3.**
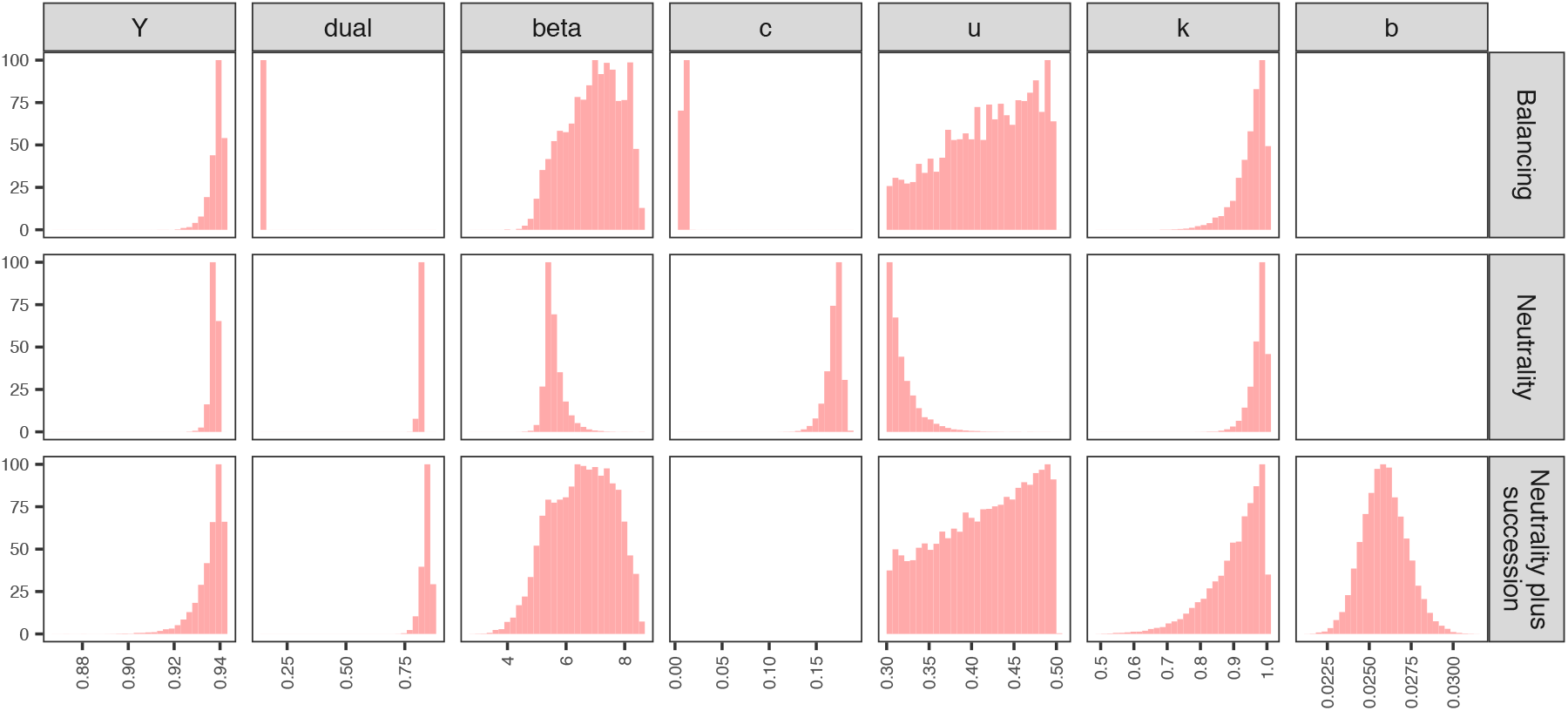
Posterior distribution for *E. coli* / penicillin, 2015. See Fig. S1 for details.

**Figure S4.**
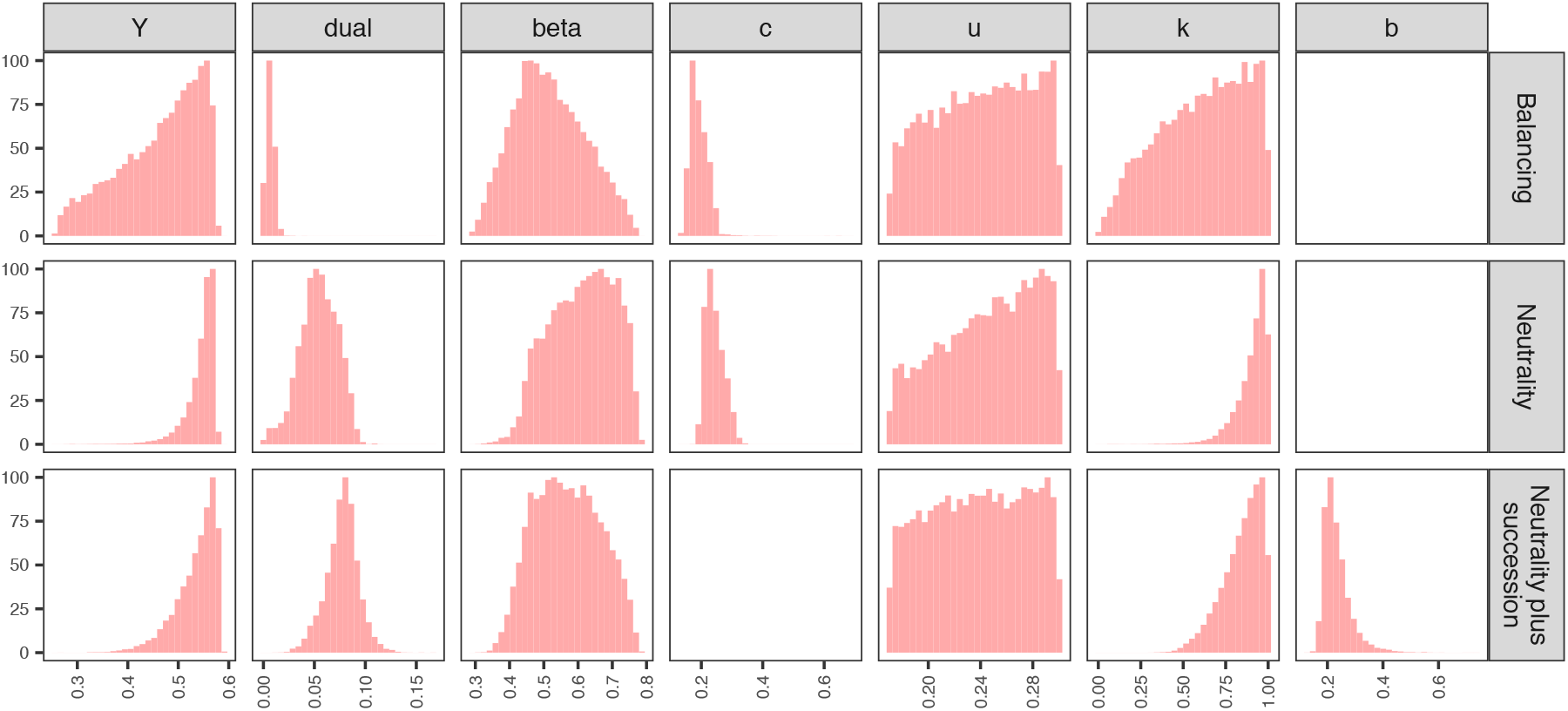
Posterior distribution for *S. aureus* / meticillin, 2015. See Fig. S1 for details.

**Figure S5.**
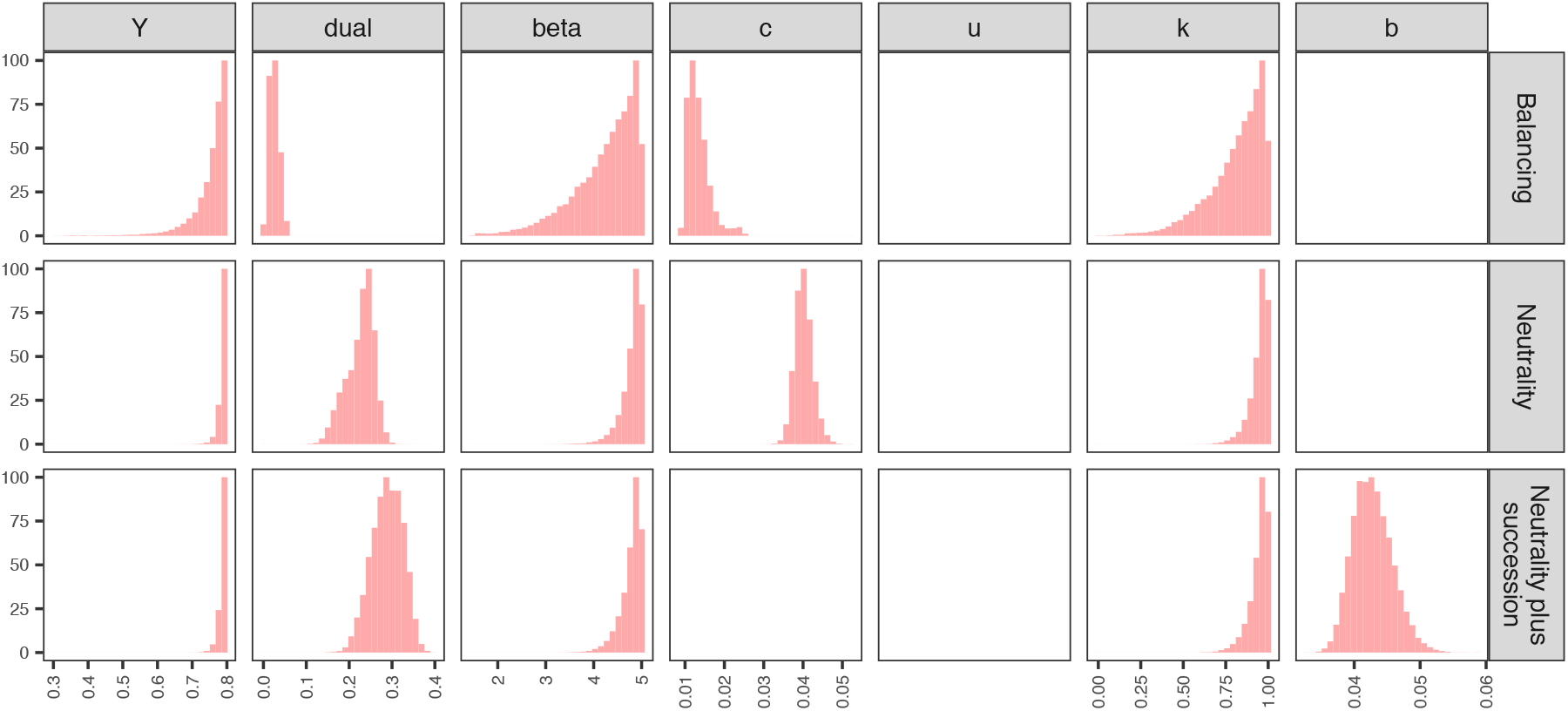
Posterior distribution for *S. pneumoniae* / macrolides, 2015. See Fig. S1 for details.

#### S2. Positive association between carriers: *k* > 1

In the main analysis, we assume that *k* measures relative efficiency of co-colonisation compared to colonisation of an uncolonised host, and hence restrict our attention to *k* ≤ 1. However, it is also possible to interpret *k* as capturing positive association between carriers, as increasing *k* increases transmission among carriers without impacting upon the overall prevalence of carriage in the population. Thus, it is possible to have *k* > 1 reflecting this mechanism. Allowing for the range 0 ≤ *k* ≤ 5, we find that (a) the fit of the within-host balancing model does not substantially improve and that (b) similar levels of coexistence are possible for the within-host neutral and within-host competition models with lower prevalence of carriage. Here, for illustration, we adopt more restrictive ranges on the prevalence of carriage: 0.2 ≤ *Y* ≤ 0.4 for *S. pneumoniae*, 0.4 ≤ *Y* ≤ 0.8 for *E. coli*, and 0.2 ≤ *Y* ≤ 0.4 for *S. aureus* (Fig. S6).

**Figure S6.**
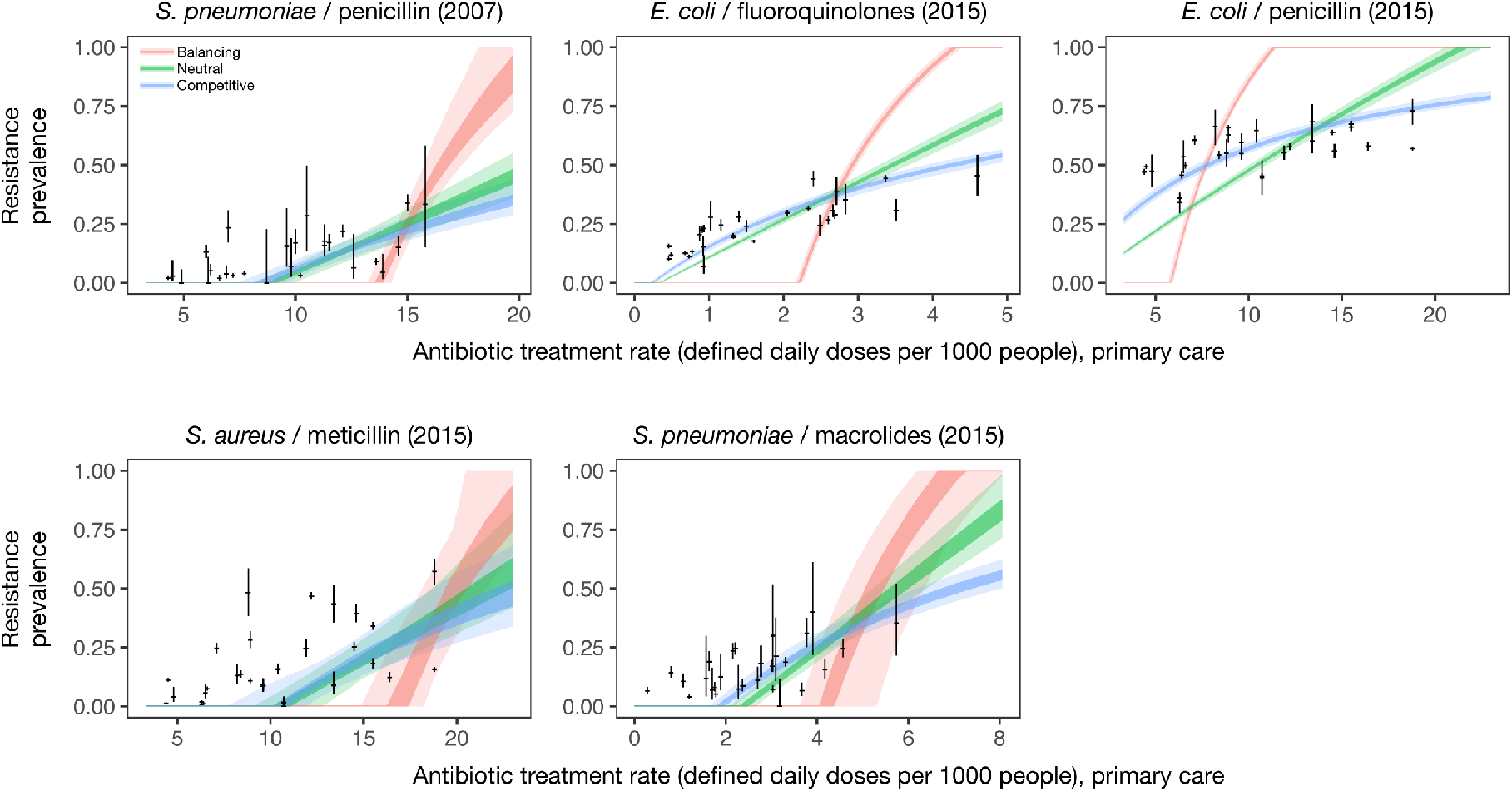
Model fits with 0 ≤ *k* ≤ 5.

##### Posterior distributions from model fitting with 0 ≤ k ≤ 5

Posterior distributions provided in Figs. S7–S11.

**Figure S7.**
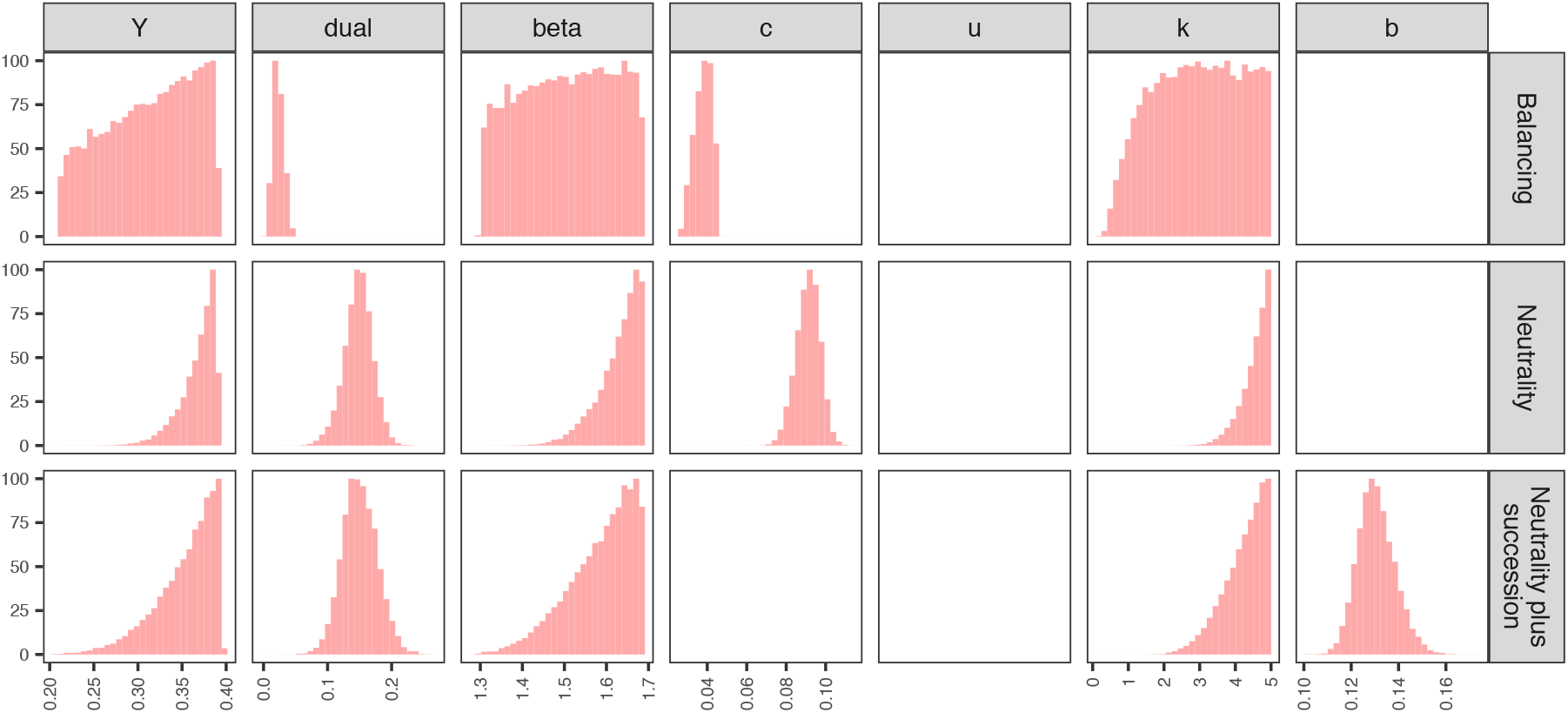
Posterior distribution for *S. pneumoniae* / penicillin (2007) data set with 0 ≤ *k* ≤ 5.

**Figure S8.**
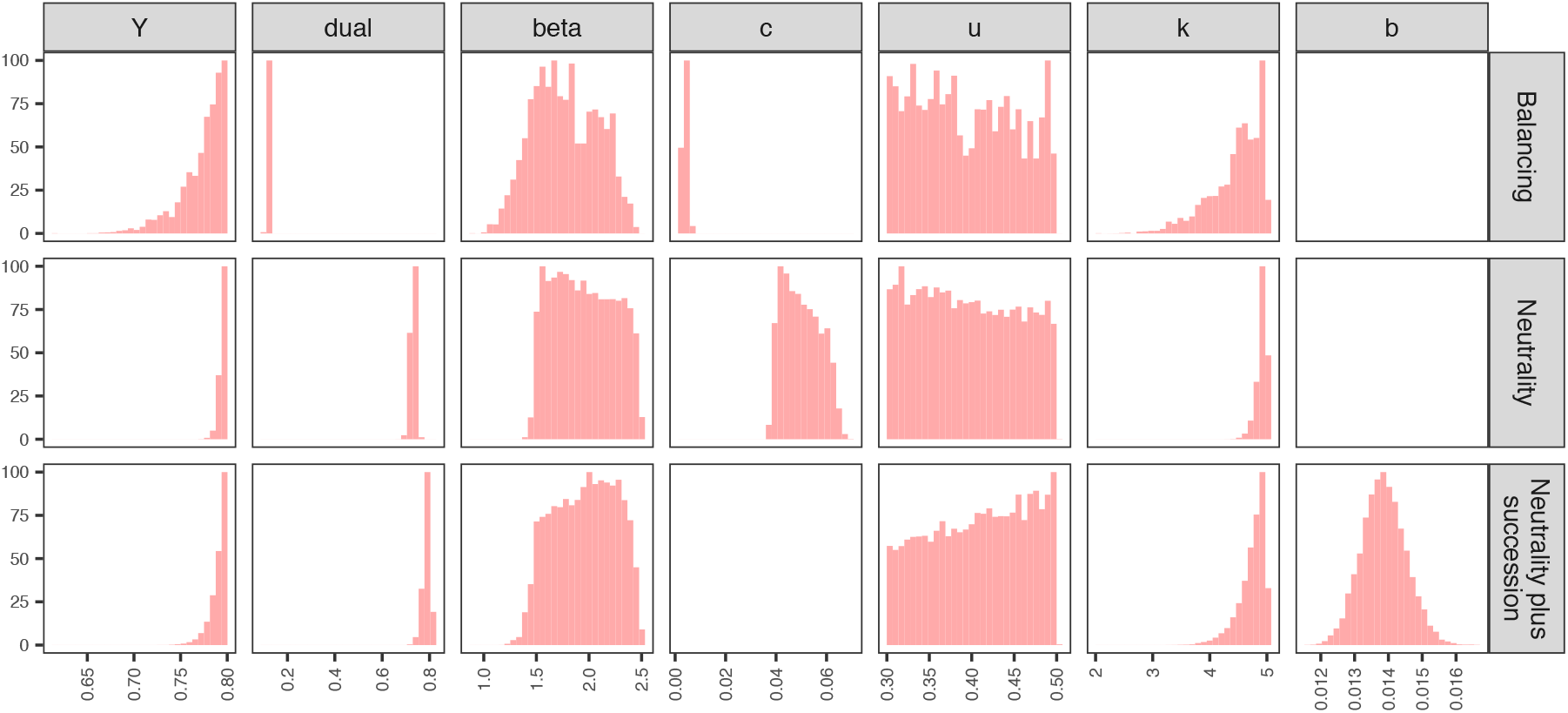
Posterior distribution for *E. coli* / fluoroquinolones (2015) data set with 0 ≤ *k* ≤ 5.

**Figure S9.**
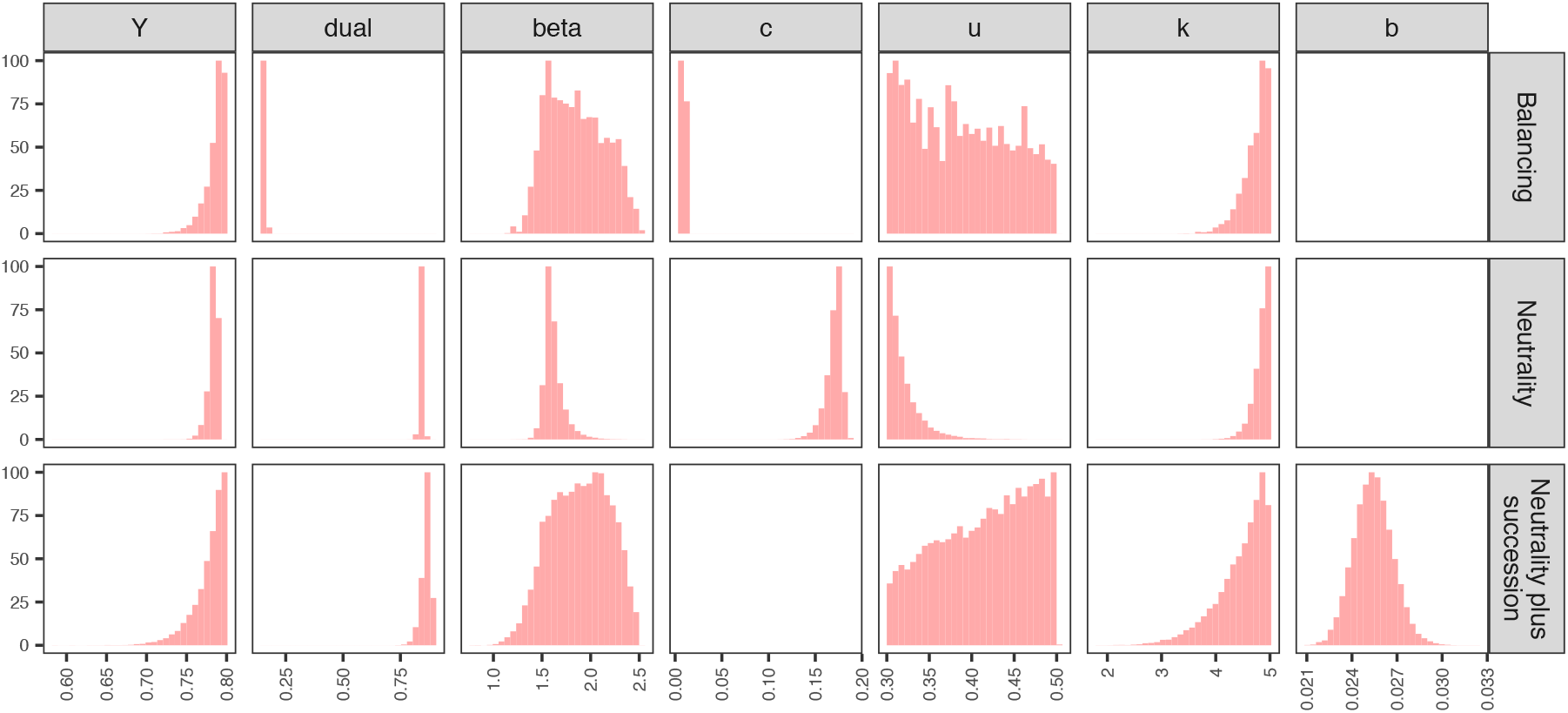
Posterior distribution for *E. coli* / penicillin (2015) data set with 0 ≤ *k* ≤ 5.

**Figure S10.**
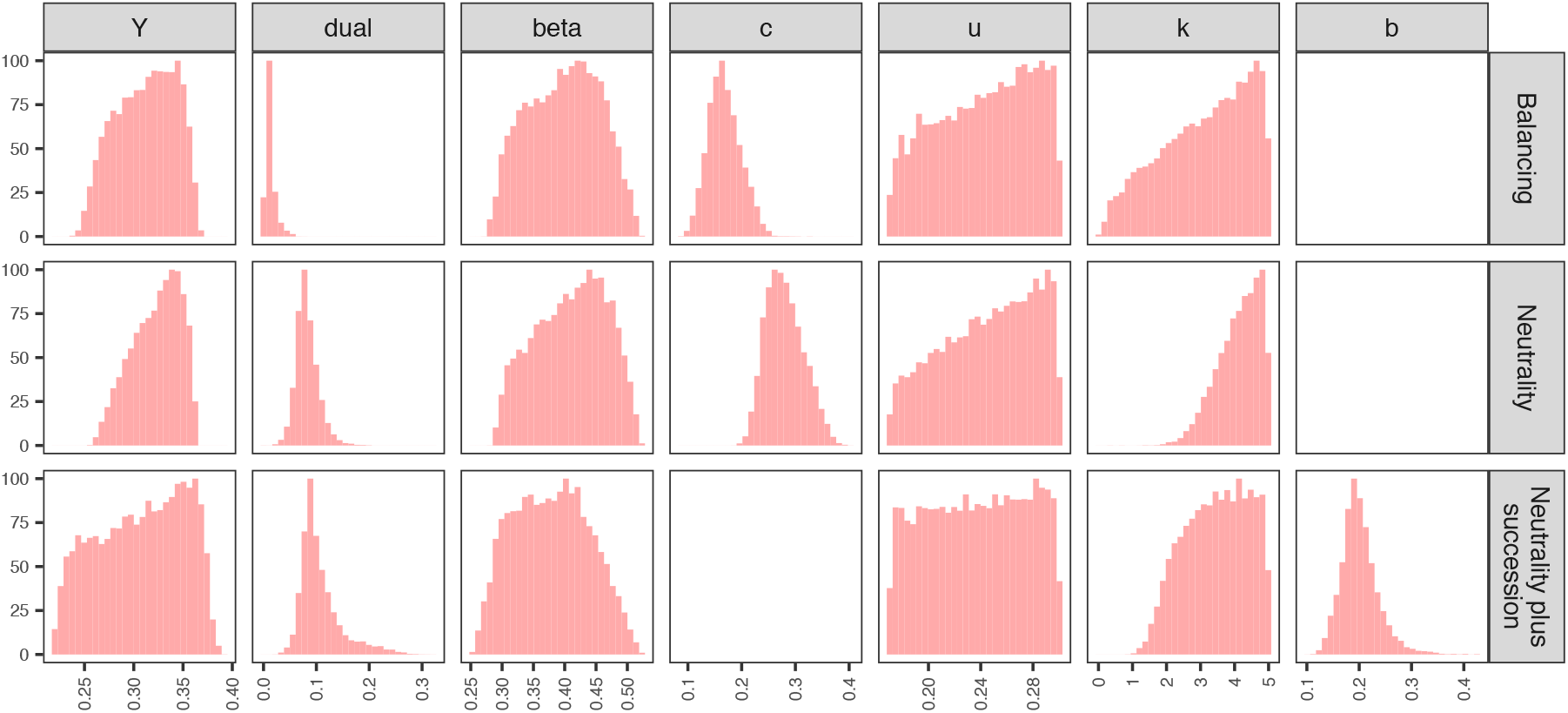
Posterior distribution for *S. aureus* / meticillin (2015) data set with 0 ≤ *k* ≤ 5.

**Figure S11.**
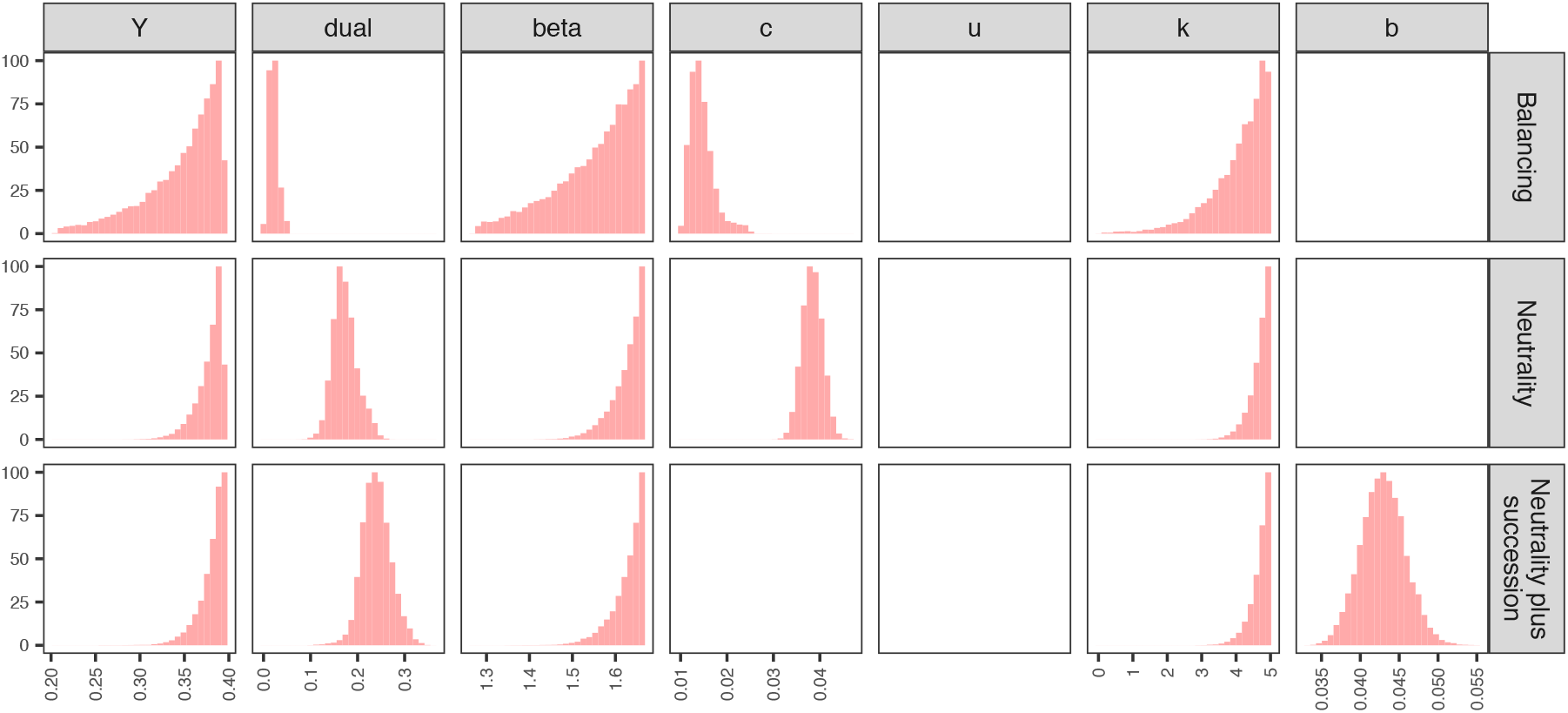
Posterior distribution for *S. pneumoniae* / macrolides (2015) data set with 0 ≤ *k* ≤ 5.

##### Relative likelihoods from model fitting with 0 ≤ k ≤ 5

We calculated the relative likelihood for the fitted models, *M_i_*, by comparison of the Akaike Information Criterion (AIC) of each model to that of the best model, M (exp[(AIC(M) - AIC(M¡)) /2]) (Table S2).

**Table S2.** Relative likelihoods for model fits with 0 ≤ *k* ≤ 5. The number of parameters, *n*, is given for each model.

| Model | Within-host<br>balancing | Within-host<br>neutrality | Within-host<br>competition |
| --- | --- | --- | --- |
| *S. pneumoniae* /<br>penicillin (2007) ($n = 3$) | $4.82 \times 10^{-5}$ | 0.453 | 1 |
| *E. coli* /<br>fluoroquinolones<br>(2015) ($n = 4$) | $1.11 \times 10^{-127}$ | $1.03 \times 10^{-12}$ | 1 |
| *E. coli* / penicillin<br>(2015) ($n = 4$) | $1.12 \times 10^{-161}$ | $1.34 \times 10^{-26}$ | 1 |
| *S. aureus* / meticillin<br>(2015) ($n = 4$) | $6.87 \times 10^{-4}$ | 0.696 | 1 |
| *S. pneumoniae* /<br>macrolides (2015) ($n = 3$) | $3.84 \times 10^{-14}$ | $1.33 \times 10^{-3}$ | 1 |

#### S3. S_R_-to-S clearance in within-host competition model

In the within-host neutral model with competition, within-host competition causes *R_S_* carriers to transition into *S_R_* carriers over time. However, an alternative model proposes that the minor complement of resistant cells in *S_R_* carriers may be outcompeted within the host over time, transitioning *S_R_* carriers to *S* carriers. To verify the robustness of our results against the clearance of resistant cells from *S_R_* carriers, we re-fit the the neutral model with within-host competition to the *S. pneumoniae* / penicillin data set assuming that *S_R_* carriers transition to *S* carriers at double the rate of the *R_S_* to *S_R_* transition, i.e. 2*b*. Adding this transition has a minimal impact on the model fit (Fig. S12). The relative likelihood of the model fitted with *S_R_*-to-S clearance is 0.926 compared to the model fitted without *S_R_*-to-S clearance.

**Figure S12.**
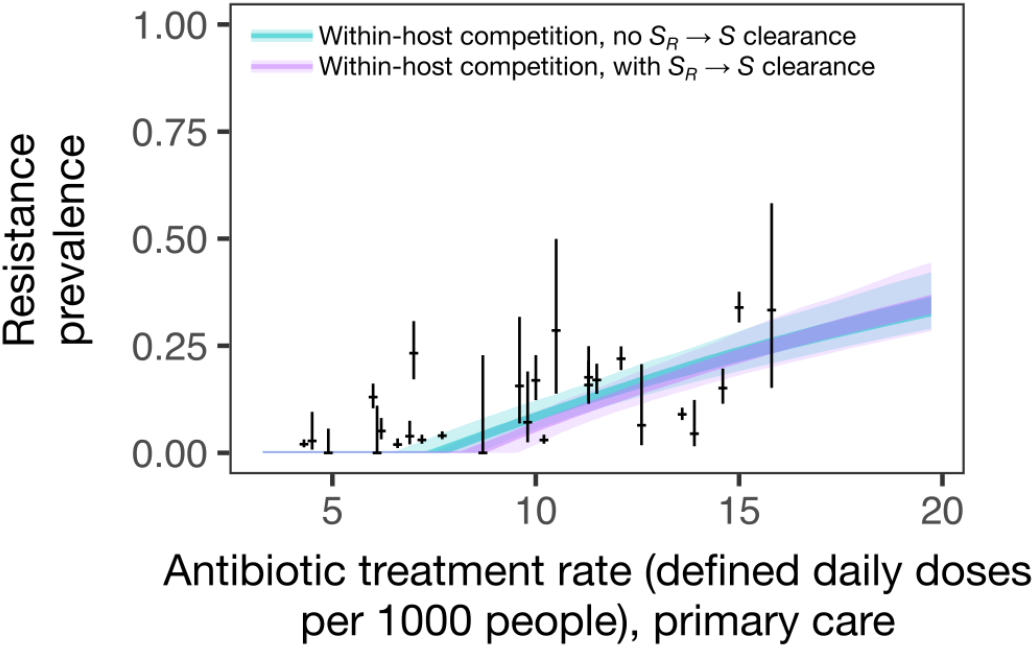
Model fits for *S. pneumoniae* / penicillin (2007) data set without and with *S_R_*-to-S clearance.

#### S4. Pneumococcal coexistence without immune memory

We found that independent clearance of serotypes alone was insufficient to support the high diversity of pneumococcal serotype carriage observed in human populations with only four serotypes maintained (Fig. S13).

**Fig. S13.**
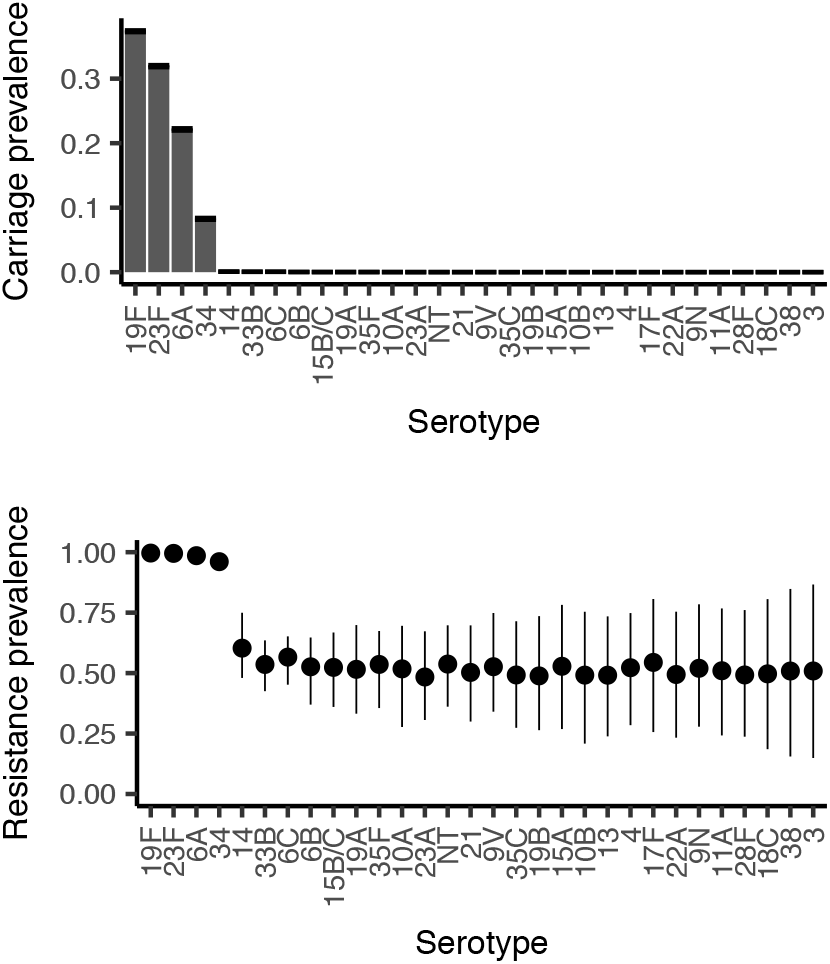
As serotypes vary greatly in duration of carriage, the mechanism of independent clearance alone is not able to reproduce observed patterns of pneumococcal carriage or resistance prevalence, as all but the four serotypes with the highest duration of carriage are eliminated. Resistance prevalence is close to 50% for eliminated serotypes as stochastic importation of strains (see *Methods*) maintains carriage of both sensitive and resistant strains of each serotype at low prevalence.

